# Reelin plasma levels predict cognitive decline in Alcohol Use Disorder: peak levels in patients expressing plasma APOE4 protein

**DOI:** 10.1101/2023.02.08.527670

**Authors:** Berta Escudero, Marta Moya, Leticia López-Valencia, Francisco Arias, Laura Orio

## Abstract

**Background:** Human apolipoprotein (APO)-E4 has been related to neuropsychiatric disorders such as Alzheimer’s disease and cognitive decline. Reelin and Clusterin share the VLDLR and ApoER2 receptors with APOE4. Here we checked the role of these components in Alcohol Use Disorder (AUD)-induced cognitive decline.

**Methods:** This is a cross-sectional study with AUD-diagnosed patients (DSM-5) (n=24) recruited from an outpatient ‘Alcohol Programme’ and matched controls (n=34). Participants were assessed by the validated ‘Test of Detection of Cognitive Impairment in Alcoholism’ (TEDCA). APOE4 presence in plasma (distinguishing APOE4 carriers and no carriers subjects) and its levels were performed by ‘e4Quant’ technique. The rest of biological markers were tested by Enzyme-Linked Immunosorbent Assay kits.

**Results:** Plasma APOE4 isoform was present in 37.5% and 58.8% of patients and controls, respectively. Quantification analyses revealed that APOE4 reached similar plasma levels in carriers independently if they were AUD subjects or controls. Circulant plasma APOE4 had a negative effect on AUD’s cognition, specifically affecting Memory/Learning (*p*<0.01, η^2^=0.15). Plasma Clusterin and Reelin increased in patients but, interestingly, Reelin plasma levels peaked in patients expressing APOE4 (*p*<0.05, η^2^=0.09), who showed reduced VLDL and ApoER2 expression in peripheral blood mononuclear cells (PBMCs). Reelin was a good predictor of cognitive loss in patients, accounting for the 42.3% and 54.0% of general intelligence and executive function impairments, respectively.

**Conclusions:** Reelin plasma levels are increased in AUD patients who express the APOE4 isoform, predicting cognitive deterioration to a great extent. Remarkably, plasma Reelin helps to differentiate between AUD patients with and without cognitive decline.

**Significance Statement:** Finding biological markers that predict a worse evolution in neuropsychiatric disorders may help to assist vulnerable patients appropriately. In this sense, in this study we found a biological marker, Reelin, which is elevated in patients with diagnosis of alcohol use disorder (AUD) that underwent an outpatient treatment. Interestingly, Reelin plasma levels were elevated in patients that also express APOE4, an aberrant protein present only in a small percentage of the population which is related to neuroinflammation and cognitive impairment (i.e. it is involved in Alzheimer’s disease). We observed that Reelin plasma levels negatively correlate with cognitive scores, being Reelin a good predictor of cognitive impairment in patients. These results may have implications for the follow-up of AUD patients in outpatient treatment.

## INTRODUCTION

Alcohol Use Disorder (AUD) is a public health issue characterized by a problematic pattern of alcohol consumption which represents a risk factor for disease, functional disability, and premature death throughout the world, generating around 3 million deaths per year (Witkiewitz et al., 2019; World Health Organization, 2014), and whose related brain damage is accompanied by a significant cognitive impairment (Visontay et al., 2021; Aharonovich et al., 2018; Wollenweber et al., 2014). The cognitive deterioration profile in AUD patients are on a spectrum (Hayes et al., 2016) and includes from lower cognitive ability or cognitive decline (Neafsey & Collins, 2011; Sabia et al., 2014) to dementia (Visontay et al., 2021). The cognitive domains with the most significant impairment appear to be visuospatial abilities, memory (working memory, learning, recall and recognition), and executive functioning (EF) (flexibility, organization, inhibition), as referred in the literature (Sachdeva et al., 2016; Hayes et al., 2016). However, difficulties have arisen from a lack of uniformity in the assessment of cognitive impairment in this pathology, without a neuropsychological test specific for these patients. At the same time, the endogenous mechanisms that could mediate cognition in this pathology are still unknown, so for this reason, we aim at offering a novel perspective using a cognitive test specifically validated in AUD (Jurado-Barba et al., 2017) that served as an instrument to evaluate the possible influence of some biomarkers on AUD-associated cognitive impairment.

One of the biomarkers widely related to cognition is the apolipoprotein (Apo)E. The human ApoE gene is polymorphic, with three alleles (ε2, ε3, ε4) encoding the different isoforms of the 35kDa lipid transport glycoprotein apolipoprotein E (APOE) (Hauser et al., 2011): APOE2, APOE3 and APOE4. The ϵ4 allele of the ApoE4 is considered dysfunctional and is expressed only in a percentage of the population (ApoE-ε4 carriers) having one or more copies (i.e. ε3/4 heterozygotes and ε4/4homozygotes). APOE-ε4 inheritance has been associated with Alzheimer’s disease (AD) (Riedel et al., 2016), being one of the most accepted genetic risk factors for developing sporadic and familial late-onset AD (Schmechel et al., 1993; Serrano-Pozo, 2021), with neuroinflammation (Duro et al., 2022; Kloske & Wilcock, 2020) and emergence of cognitive decline (Reivang et al., 2010; Montagne et al., 2020). While the influence of ApoE-ε4 in AD (in which there is a progressive loss of cognitive ability, particularly in memory) has been extensively characterized, little is known about its role in AUD. Some prospective studies in middle-aged and older adults showed that the ApoE-ε4 genotype may moderate the relationship between general cognition and alcohol consumption (Slayday et al., 2020; Downer et al., 2014), so that ApoE-ε4 carriers have an increased vulnerability to alcohol-induced neurotoxicity, indirectly influencing cognition (Kim et al., 20212). In another study with older adults (from 70 to 90 years old), ApoE-ε4 carriers were more likely to develop dementia, but no significant interaction was found with alcohol consumption (Heffernan et al., 2016). To our knowledge, the studies performed to date related to alcohol consumption did not consider subjects with clinical diagnosis of AUD, they are genetic, with no measurement or APOE4 plasma levels, and did not check the impact of peripheral APOE4 on AUD-associated cognitive impairments.

The APOE signaling pathway involves APOE binding to lipoprotein-receptors: the very-low density liporeceptor (VLDLR) and the apolipoprotein E receptor 2 (ApoER2) (Lane-Donovan & Herz, 2017), both members of the LDL receptor gene family. This pathway also involves the signaling molecule Reelin (Dlugosz & Nimpf, 2018), a large 420-kDa glycoprotein of the extracellular matrix predominantly expressed by Cajal-Retzius (CR) cells (reviewed in Causeret et al., 2021) that plays a pivotal role in brain development and adult brain function by being involved in neuronal migration, development of lamellar brain structures and synaptic plasticity of the cerebral cortex (D’Arcangelo et al., 1995; Ogawa et al., 1995, Alcántara et al., 1998; Herzet al., 2006; Tissir & Goffinet, 2003). While ApoE-ε4 has been associated with cognitive impairment and loss, Reelin, sharing the same receptors as APOE4 (Dlugosz and Nimpf, 2018), has been proposed to improve cognition and reduce memory impairment (Stranahan et al., 2013), being considered in the scientific community as a marker of protection against cognitive deterioration. In fact, Reelin gene haploinsufficiency has been linked to cognitive impairment (Ishii et al., 2016).

As APOE and Reelin, there is another apolipoprotein, the Apolipoprotein J (APOJ) or Clusterin, which is also a ligand for VLDLR and ApoER2 (Dlugosz & Nimpf, 2018; Leeb et al., 2014). APOJ is a multifunctional heterodimericglycosylated protein of 75-80 kDa, composed of two 40 kDa subunits (named α- and β-chains) (de Silva et al., 1990), whose plasma levels have been also related to changes in cognition. Like its partner APOE4 apolipoprotein, APOJ has been linked to the progression of AD and brain atrophy in mild cognitive impairment (Thambisetty et al., 2012), worse cognitive scores in Diabetes Mellitus and AD (Ha et al., 2020), and dementia (Yang et al., 2019).

Despite some evidence pointing out that these three molecules (APOE4, Reelin and Clusterin) with common receptors may modulate cognitive performance in a variety of neuropsychological disorders, no studies have been conducted about their possible contribution in patients suffering from AUD, where cognitive impairment is prominent. Our present study explored the specific contribution of peripheral APOE4 (presence or absence of plasma APOE4 and concentration of APOE4 in carriers of both groups) in a cohort of abstinent AUD-diagnosed patients and control subjects and studied its effect on cognitive function in detail by checking specific cognitive domains. We hypothesized that those patients who express APOE4 in plasma will show poorer cognition than those who do not express plasma APOE4. We investigated the affectation of different cognitive domains. In addition, we checked the role of plasma levels of Reelin and Clusterin in AUD’s cognition and explored the differences with controls according to the presence of APOE4 and sex. We also described the presence of VLDLR and ApoER2 in peripheral blood mononuclear cells (PBMCs) and their modulation in patients. Results indicate that the peripheral Reelin/APOE4 pathway may be altered in AUD patients with worse cognitive performance.

## METHODS

### Study Participants

A total of 76 subjects (white Caucasian) were recruited (see Flowchart, Fig. 1) and divided into two groups. (1) AUD group: 39 abstinent patients, recruited from the*Hospital Universitario 12 de Octubre* (Madrid, Spain) (outpatient ‘Alcohol Programme’; see Supplement 1) and (2) control group: 37 healthy subjects with no history of drug abuse.

**FIGURE 1.**
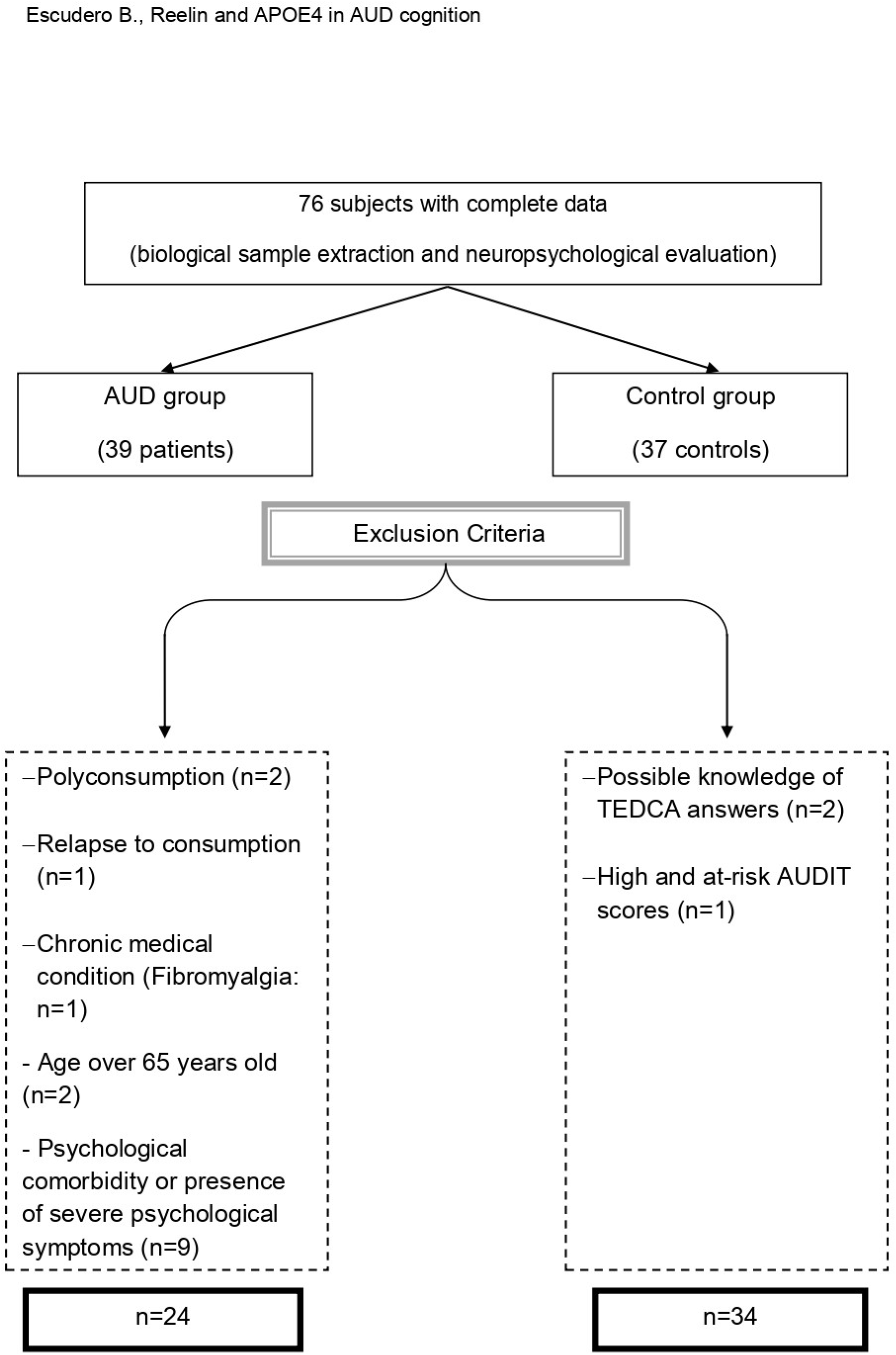
Flowchart on sample recruitment. Participants were divided into two experimental groups: AUD group and control group. Based on the exclusion criteria previously established for the study, the final selected sample is established. Abbreviations. AUD: Alcohol Use Disorder; TEDCA: Test of detection of cognitive impairment in alcoholism; AUDIT: Alcohol Use Disorders Identification Test.

#### Inclusion criteria

Age between 18 and 65 years old. In case of patients, AUD diagnosis based on DSM-5 criteria (American Psychiatric Association, 2014) and alcohol abstinence for at least 4 weeks before testing.

#### Exclusion criteria

History of abuse or dependence to other drugs (including alcohol in the control group), except tobacco; any psychiatric comorbidity or concomitant psychological disorder or presence of severe psychological symptoms; chronic medical condition, presence of infectious diseases (HIV infection and/or hepatitis) and chronic use of anti-inflammatory medication.

After corroborating the eligibility to the study, 24 patients (17 men and 7 women) were included in the AUD group and 34 (16 men and 18 women) in the control group.

This study was approved by the Research Ethics Committee of the *Hospital 12 de Octubre, Madrid* (Spain), Nº CEIm: 19/002, and was conformed to the provisions of the World Medical Association Declaration of Helsinki. Each participant signed a written informed consent individually and all data were coded to maintain anonymity and confidentiality.

### Clinical assessment

The diagnosis of alcohol disorder in patients were examined based on DSM-5 criteria (American Psychiatric Association, 2014), whereas in controls, risky alcohol consumption was assessed using ‘Alcohol Use Disorders Identification Test’ (AUDIT) (Babor et al., 2001). Comorbidity in patients and controls was controlled using the Spanish version of the ‘Mini International Neuropsychiatric Interview’ (MINI) (Ferrando et al., 1998) and under psychiatrists’ medical criteria (only in patients) (see Supplement 2).

For both groups, a semi-structured interview was also applied on alcohol use history and use or abuse of other drugs, including tobacco (Polyconsumption). Participants were asked whether they had smoked regularly for at least 1 year (no, yes and former) (Table S2 Supplement 2). In addition, Beck Depression Inventory-II (BDI-II) (Beck et al., 1996; Sanz et al., 2003) and Beck Anxiety Inventory (BAI) (Beck & Steer, 2011) were administered to patients and controls to assess depression and anxiety symptoms, respectively.

### Neuropsychological testing

Participants were assessed by TEDCA (“Test of detection of cognitive impairment in alcoholism”) (Jurado-Barba et al., 2017), a test that has been specifically validated in the alcohol-dependent patient population and presents adequate psychometric properties. TEDCA is used in the hospital clinical practice as a screening test that provides a snapshot of cognitive functioning (General Cognitive Functioning (GCF)) based on a compendium of 7 tests that assess three specific cognitive functions: Visuospatial Cognition, Memory/Learning and EF (Supplement 3). The final score meaning general intelligence or GCF results from the sum of the three previous ones. The maximum score to be obtained in TEDCA is 18 for general intelligence and 6 points for each cognitive domain. Impairment is established at ≤10.5 (test cut-off point) for general intelligence and at ≤ 3 (cut-off point) for specific cognitive domains. Therefore, subjects with equal or inferior scores to the cut-off points exhibit cognitive decline.

### Sample collection and processing

Blood extractions were performed in the morning (8:00-10:00 a.m.) after 8-12 h of fasting and extracted into 10mL BD vacutainer® tubes with K2-EDTA anticoagulant (BD, Franklin Lakes, NJ, USA). Blood samples were immediately processed to obtain plasma and PBMCs based on our previous studies (Orio et al., 2018). Briefly, plasma was extracted by blood centrifugation at 1800 rpm for 10 minutes at 4ºC and PBMCs extraction process consisted of the layer-by-layer separation of different blood components using the Ficoll technique, producing a thin layer of lymphocyte cells that were reabsorbed. Plasma/PBMCs were coded and stored at -80ºC until further use.

### APOE4, Reelin, Clusterin, VLDLR and ApoER2 detection

APOE4 expression in plasma is only present in a percentage of individuals carrying the ε4 allele, either ApoE-ε3/4 (heterozygous) or ApoE-ε4/4 (homozygous) gene carriers. Since the presence of the heterozygous or homozygous ε4 allele may result in different total APOE4 plasma levels (Fukumoto et al. 2003), we directly measured the protein levels of the APOE4 isoform in plasma by the e4Quant patented technique (Biocross S.L.; patent nº EP15382537.7), using the chemistry analyzer KROMA-PLUS (Linear Chemicals SU, Barcelona) (Calero et al., 2018). e4Quant is a quantitative test and a validated method based in turbidimetry that detect and quantify specifically APOE4 and not other isoforms (i.e. APOE3 or APOE2) in human plasma, which has been used as a predictor of AD progression (Calero et al., 2018). e4Quant takes advantage of the capacity of APOE4 to bind to polystyrene surfaces as capture and a specific anti-APOE4 monoclonal antibody as reporter.

Reelin, Clusterin (in plasma), VLDLR and ApoER2 (in PBMCs) were determined by commercially available Enzyme-Linked Immunosorbent Assay (ELISA) kits following the manufacturer’s instructions (Supplement 4).

### Statistical analyses

Comparisons between patients and controls were analyzed by Fisher’s exact test, Chi-Square test or Student’s t-test, according to the statistical assumptions for each test, and shown as number and percentage of subjects [N (%)] or mean and standard error of the mean [mean (±SEM)].

Analyses of covariance (ANCOVA) were performed considering the assumptions of covariate independence and homoscedasticity by Levene’s test, including age as covariate. ANCOVAs are shown as follows: F-statistic (factor/error degrees of freedom), *p* value and effect size partial eta-squared η^2^. Bonferroni *post hoc* test was run when appropriate. Figures are expressed as scatterplots, with mean ± SEM.

The relationship between biomarkers and patient’s cognition was studied using Spearman’s rank coefficients (r) (adjusted R^2^). To account for multiple testing in the correlations, we adjusted the p-values by controlling the false discovery rate (FDR) at 5% using the Benjamini-Hochberg procedure. This type of correction allows restricting the occurrence of false-positive findings among all nominal significant findings to a certain threshold.

Patients were divided into those with and without cognitive impairment in each domain and Binary Logistic Regression was performed for Reelin (the only biomarker who showed differences in cognition in the mean difference analysis (Student’s t-test)). Reelin was transformed to standard Z scores for a better interpretation of the odds ratios, because the transformation did not affect the *Wald* values and corresponding *p* values. A Receiving Operating Characteristic (ROC) curve analysis was performed for GCF and EF to determine the thresholds beyond which Reelin could be regarded as a risk factor. The identification of risk factors (APOE4, Reelin, Clusterin, VLDLR and ApoER2) for the prognosis of cognitive performance was also assessed using a Linear Regression model.

Statistical analyses were performed through IBM SPSS Statistical Version 25.0 software (IBM, Armonk, USA) and GraphPad Prism^©^ version 8.00. (GraphPad Software, Inc., CA, USA). A *p*-value <0.05 was considered statistically significant.

## RESULTS

### Sample demographics and neuropsychological findings

Socio-demographic comparisons between patients and controls are presented in Table 1. The groups were matched on all potential contaminant variables except for age (*p*<0.01), which was adjusted as covariate in the ANCOVAs analyses. For both patients and controls, BMI was in the normal range, there were healthy cholesterol levels and no difference in the number of current smokers. The most used psychiatric medication in patients was disulfiram, followed by antidepressants (no one in the control group received psychiatric medication).

**Table 1.**
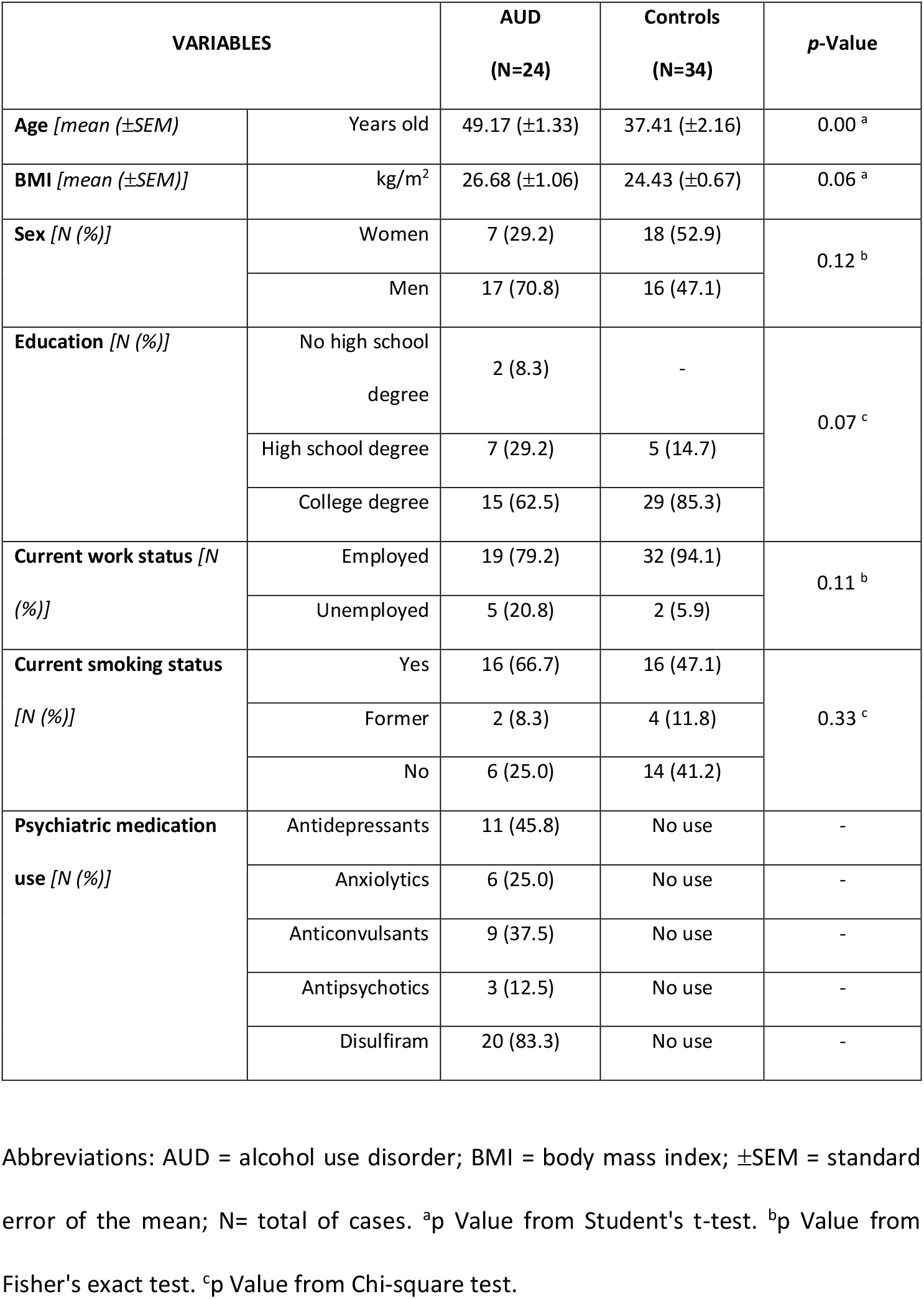
Socio-demographic and pharmacological variables between AUD and Controls

| VARIABLES |  | AUD<br>(N=24) | Controls<br>(N=34) | p-Value |
| --- | --- | --- | --- | --- |
| Age [mean (±SEM)] | Years old | 49.17 (±1.33) | 37.41 (±2.16) | 0.00 <sup>a</sup> |
| BMI [mean (±SEM)] | kg/m<sup>2</sup> | 26.68 (±1.06) | 24.43 (±0.67) | 0.06 <sup>a</sup> |
| Sex [N (%)] | Women | 7 (29.2) | 18 (52.9) | 0.12 <sup>b</sup> |
|  | Men | 17 (70.8) | 16 (47.1) |  |
| Education [N (%)] | No high school degree | 2 (8.3) | - | 0.07 <sup>c</sup> |
|  | High school degree | 7 (29.2) | 5 (14.7) |  |
|  | College degree | 15 (62.5) | 29 (85.3) |  |
| Current work status [N (%)] | Employed | 19 (79.2) | 32 (94.1) | 0.11 <sup>b</sup> |
|  | Unemployed | 5 (20.8) | 2 (5.9) |  |
| Current smoking status [N (%)] | Yes | 16 (66.7) | 16 (47.1) | 0.33 <sup>c</sup> |
|  | Former | 2 (8.3) | 4 (11.8) |  |
|  | No | 6 (25.0) | 14 (41.2) |  |
| Psychiatric medication use [N (%)] | Antidepressants | 11 (45.8) | No use | - |
|  | Anxiolytics | 6 (25.0) | No use | - |
|  | Anticonvulsants | 9 (37.5) | No use | - |
|  | Antipsychotics | 3 (12.5) | No use | - |
|  | Disulfiram | 20 (83.3) | No use | - |

Abbreviations: AUD = alcohol use disorder; BMI = body mass index; ± SEM = standard error of the mean; N= total of cases. <sup>a</sup>p Value from Student's t-test. <sup>b</sup>p Value from Fisher's exact test. <sup>c</sup>p Value from Chi-square test.

Cognitive data from AUD patients and controls are presented in Table 2. Patients showed significant lower cognitive performance for all domains. We found that 79.2% of patients presented Memory/Learning deficits, 41.7% EF alterations, and 12.5% Visuospatial deficits, fulfilling the 29.2% of all of them a GCF deficit. The Memory/Learning domain was the most affected domain. It is to note that although the mean in patients was at the cut-off limit to consider impairment, most patients (80%) presented Memory/Learning impairment (scores < 3.0) being the total count compensated by patients with no impairment in this subdomain. The most preserved domain was Visuospatial ability, with relatively high scores, which may have masked the final number of patients with cognitive impairment (GCF).

**Table 2.** Cognitive data and differences between AUD patients and controls

| Cognitive domains |  | GCF | Visuospatial Cognition | Memory /Learning | EF |
| --- | --- | --- | --- | --- | --- |
| Direct scores<br>[mean (±SEM)] | AUD<br>(n=24) | 11.75 (±0.67) | 4.62 (±0.27) | 3.04 (±0.30) | 4.04 (±0.30) |
| | Controls<br>(n=34) | 14.85 (±0.33) | 5.38 (±0.13) | 4.74 (±0.16) | 4.74 (±0.19) |
| p-Value |  | 0.00 <sup>a</sup> | 0.01 <sup>a</sup> | 0.00 <sup>a</sup> | 0.04 <sup>a</sup> |

Abbreviations. ± SEM: standard error of the mean. n= total of cases. AUD: alcohol use disorder. GCF: General Cognitive Functioning. EF: Executive functioning. <sup>a</sup>p Value from Student's t-test.

### Status of markers in AUD pathology

Biochemical results are presented in Table 3. The patented eQuant technique showed the number and percentage of patients and controls expressing plasma APOE4 and the quantitative results of APOE4 plasma levels for those subjects [mean (±SEM)].

**Table 3.** APOE4, Reelin, Clusterin, VLDLR and ApoER2 results

| VARIABLES |  | AUD (N=24) | Controls (N=34) | p-value |
| --- | --- | --- | --- | --- |
| **Plasma APOE4 expression**<br>[N (%)] | Yes | 9 (37.50) | 20 (58.80) | 0.11 <sup>a</sup> |
|  | No | 15 (62.50) | 14 (41.20) |  |
| **APOE4 levels (ApoE4-ε4 carriers)**<br>(μg/mL) [mean (±SEM)] in those expressing plasma APOE4 (0 values are not considered) |  | 11.28 (±3.10) | 11.08 (±1.43) | 0.95 <sup>b</sup> |
| **Reelin levels** (ng/mL) [mean (±SEM)] |  | 0.42 (±0.41) | 0.30 (±0.27) | 0.02 <sup>b</sup> |
| **Clusterin levels** (μg/mL) [mean (±SEM)] |  | 46.65 (±4.67) | 30.72 (±3.58) | 0.01 <sup>b</sup> |
| **VLDLR levels** (ng/mL total protein) [mean (±SEM)] |  | 4.14 (±0.56) | 5.32 (±0.60) | 0.17 <sup>b</sup> |
| **ApoER2 levels** (ng/mL total protein) [mean (±SEM)] (*) |  | 3.36 (±0.48) | 3.20 (±0.35) | 0.78 <sup>b</sup> |

Abbreviations: ±SEM = standard error of the mean. n= total of cases. (\*) means that the n=23 due to the presence of an outlier in group AUD. <sup>a</sup>p Value from Chi-square test, <sup>b</sup>p Value from Student's t-test.

Contrary to expectations and in our cohort, there was a higher percentage of subjects expressing plasma APOE4 in controls (58.80%) versus patients (37.50%), but without significant differences (p=0.11, Chi square test) (Table 3). As said before, APOE4 expression in plasma is only present in a percentage of individuals carrying the ε4 allele, either ApoE-ε3/4 (heterozygous) or ApoE-ε4/4 (homozygous) gene carriers. Thus, the percentage of carriers found in controls was higher than expected. To investigate if there were differences in the amount of APOE4 expressed in plasma in carriers, we compared the levels of plasma APOE4 considering only the subjects that express the apolipoprotein in each experimental group. We observed that the amount of APOE4 in plasma reached similar levels in both experimental groups (p=0.95 Student’s t test). Thus, although we randomly recruited more APOE4 carriers in the control group, plasma APOE4 levels reached in carrier controls were similar to those of carrier patients (Table 3).

Regarding Reelin levels (Table 3), a 3-way ANCOVA controlling for age showed a main effect of *group* [F (1,50)=7.65, *p*=0.008, η^2^=0.13] (with higher levels in patients), a main effect of *APOE4* presence [F (1,50)= 5.06, *p*=0.029, η^2^=0.09] (with increased levels in ApoE-ε4 carriers), and an interaction between *group*APOE4* [F (1,50)=4.75, *p*=0.034, η^2^=0.09] (Fig. 2A). *Post hoc* analyses revealed that there was a difference between Reelin levels in AUD patients expressing and not expressing the APOE4, which did not appear in controls, so that Reelin levels were favored to increase when the patient was an ApoE-ε4 carrier (see Fig. 2A). There was no effect of *sex* or its interaction with *group* or *APOE4* (*p*>0.05; data not shown).

**FIGURE 2.**
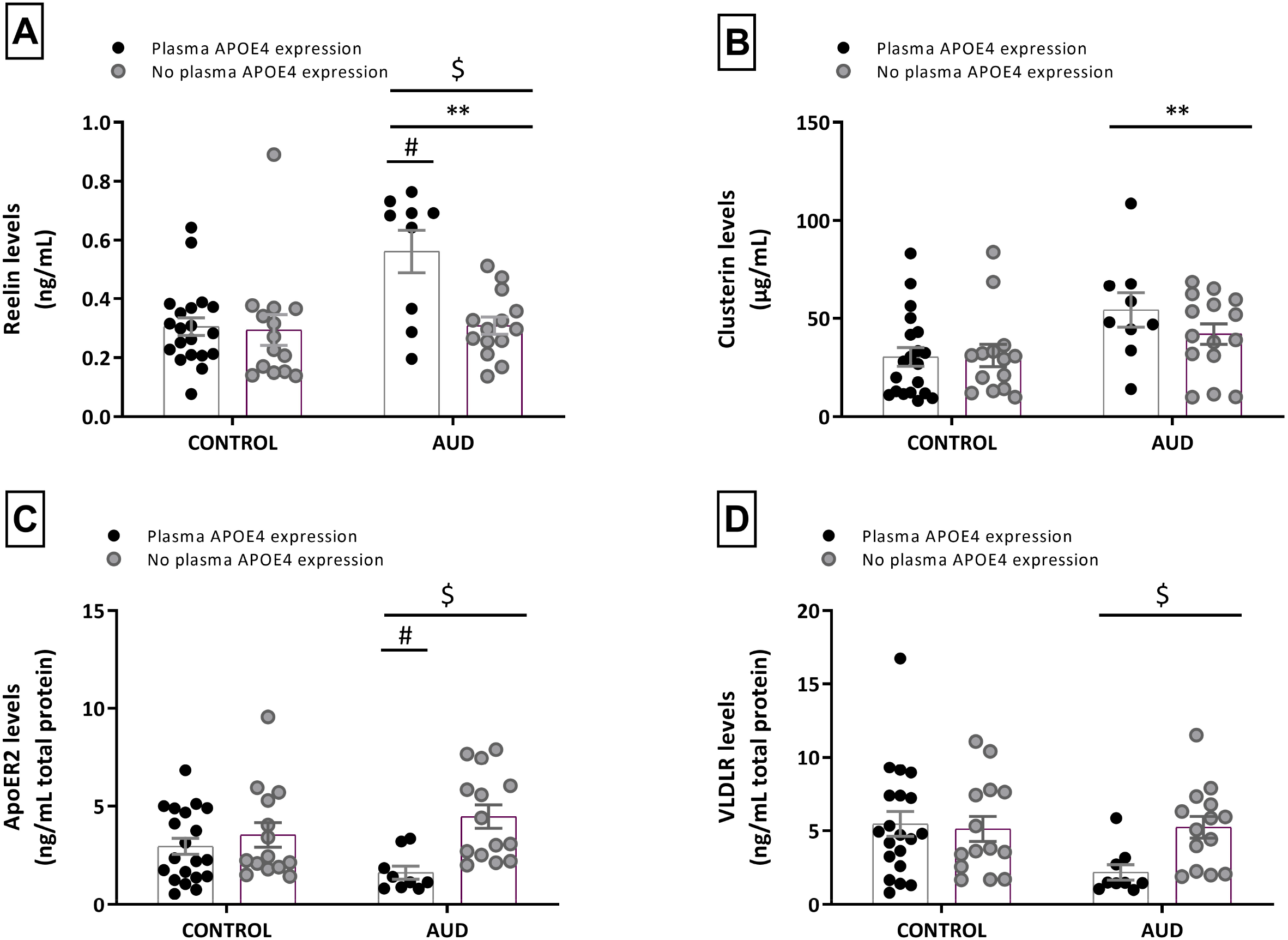
Levels of (**A**) Reelin, (**B**) Clusterin, (**C**) ApoER2 and (**D**) VLDLR in AUD patients and controls by APOE4, group and sex. Results are presented as mean ± SEM. Three-way ANCOVA, controlling for age. Overall effect of group: “**” p<0.01; APOE4: “#” p<0.05 and analysis of the interaction (group x APOE4): “$” p<0.05. Bonferroni *Post hoc* test after interaction.

Clusterin levels (Table 3) analyses revealed a *group* effect [F (1,50)=7.79, *p*=0.007, η^2^=0.13] (with higher levels in patients) and no effect of *sex, APOE4* or interactions (3-way ANCOVA controlling for age; *p*>0.05; data not shown). In this case, the condition of being an APOE4 carrier or not did not significantly influence the levels of Clusterin (although a trend could be observed) (Fig. 2B).

We detected presence of the receptors ApoER2 and VLDLR in PBMCs in patients and controls (Table 3). Regarding ApoER2, a 3-way ANCOVA controlling for age for ApoER2 revealed a main effect of *APOE4* [F (1,48)=8.69, *p=*0.005, η^2^=0.15] (decreased levels in APOE4 carriers) and an interaction between *group*APOE4* [F (1,48)=5.03, *p*=0.029, η^2^=0.09] (Fig. 2C). *Post hoc* analyses revealed that there was a difference between APOE4 carriers vs. non-carriers in patients which did not appear in controls, so that ApoER2 PBMCs levels may decrease in patients expressing APOE4 plasma levels. There was no effect of *sex* or its interaction with *group* or *APOE4* (*p*>0.05; data not shown). Regarding VLDLR, there was a main effect of *group*APOE4* [F (1,49)=4.81, *p=*0.033, η^2^=0.09] (Fig. 2D), with the lowest levels in AUD patients carrying APOE4.

### Effect of markers on cognition in alcohol-dependent patients

#### Effect of the presence of plasma APOE4 (carriers/non-carriers)on cognition

ANCOVA analyses controlling for age showed that: (a) all cognitive domains were significantly affected by *group* (lower scores in patients, as expected) (Fig. 3 A,B,C,D) [GCF: F (1,53)=18.20, *p*=0.000, η^2^=0.26; Visuospatial Cognition: F (1,53)=4.24, *p*=0.044, η^2^=0.07; Memory/Learning: F (1,53)=24.26, *p*=0.000, η^2^=0.31; EF: F (1,53)=5.86, *p*=0.019, η^2^=0.10]; (b) GCF, Memory/Learning and EF were influenced by the expression of plasma *APOE4* (as indicative of ApoE-ε4 carriers) (lower scores in carriers) (Fig. 3 A,C,D) [GCF: F (1,53)=7.24, *p*=0.009, η^2^=0.12; Memory/Learning: F (1,53)=9.17, *p*=0.004, η^2^=0.15; EF: F (1,53)= 4.12, *p*=0.047, η^2^=0.07. No effect in Visuospatial Cognition: F (1,53)=0.36,*p*>0.05, η^2^=0.01]; (c) Memory/Learning was influenced by the interaction between *group*APOE4 presence* (Fig. 3C) [F (1,53)=4.81, *p*=0.033, η^2^=0.08]. *Post hoc* analyses identified a difference in Memory/Learning performance between AUD patients who expressed APOE4 versus those who didn’t express APOE4 that was not found in the control group, with lower memory performance in those expressing APOE4. Based on these results, presenting APOE4 favors a detrimental effect on general cognitive performance (except for Visuospatial Cognition) with a clear trend in patients. The intrinsic detrimental effect of AUD diagnosis is significantly more pronounced for Memory/Learning when dealing with a patient expressing plasma APOE4. Taking into consideration all four domains (Fig. 3 A,B,C,D), the lowest performance was found for the Memory/Learning domain, precisely in these patients expressing APOE4.

**FIGURE 3.**
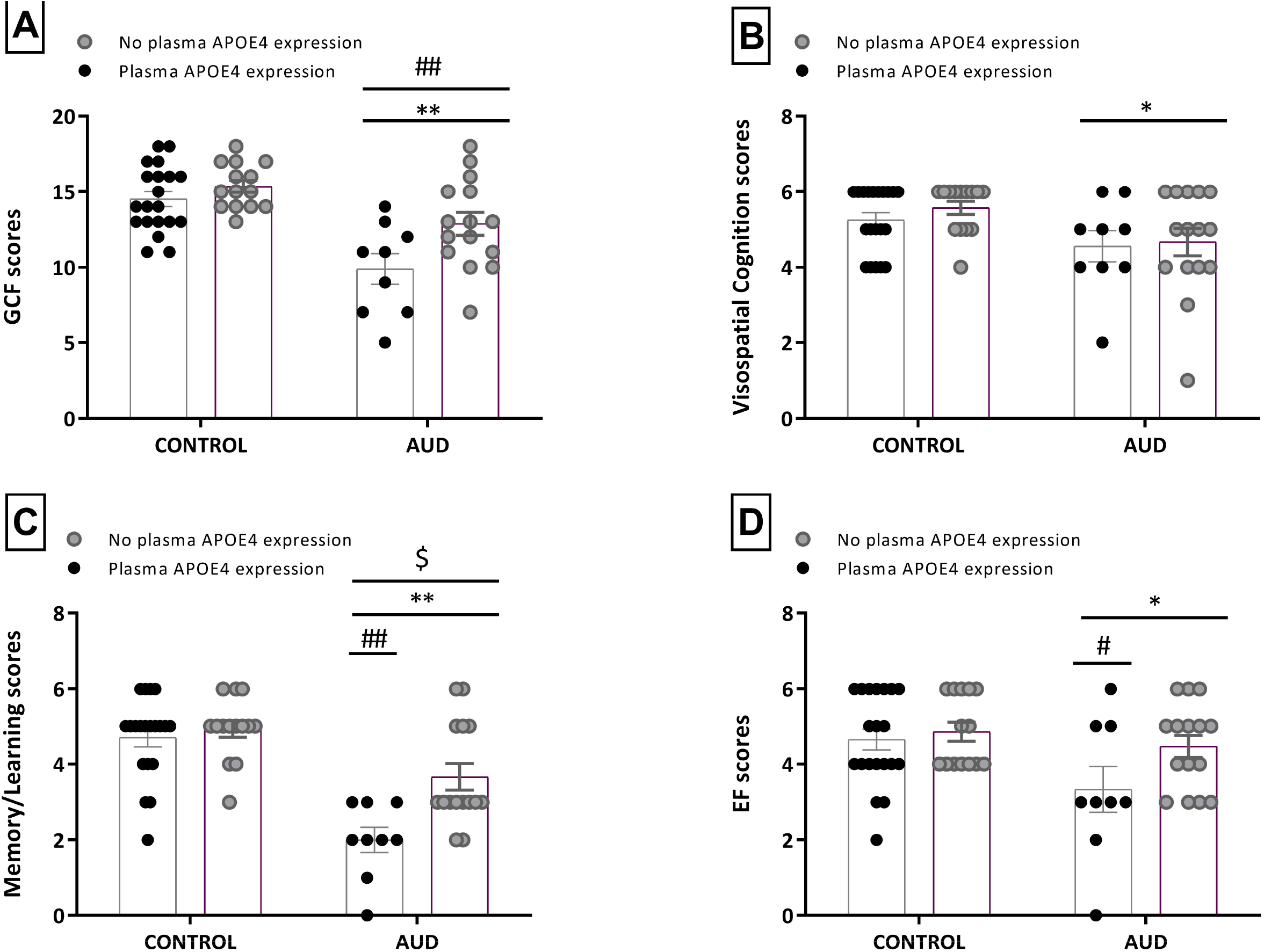
Effect of ApoE-ε4 on cognitive performance in patients and controls. Two-way ANCOVA, controlling for age. (**A**) GCF: General Cognitive Functioning (**B**) Visuospatial Cognition (**C**) Memory/Learning (**D**) EF: Executive Functioning. Results as mean ± SEM. “**” p<0.01 and “*"p<0.05 denotes a main effect of group; “##” p<0.01 and “#” p<0.05 denotes a main effect of APOE4. Analysis of the interaction (group*APOE4) in (C):"$” p<0.05. Bonferroni *post hoc* test after interaction.

#### Plasma levels of APOE4, Reelin and Clusterin and cognition

Correlation analyses between cognition scores and biomarkers (APOE4, Reelin and Clusterin), disregarding the experimental groups, were only significant for Reelin in GCF, Memory/Learning and EF [GCF: r=-0.50, *p*<0.01; Memory/Learning: r=-0.53, *p*<0.01; EF: r=-0.43, *p*<0.01]. There was no significant correlation for APOE4 or Clusterin (*p*>0.05; data not shown). Interestingly, when both groups were analyzed separately, Reelin only showed significance in patients (and not in controls; GCF, Memory/Learning and EF [GCF: r=-0.74, *p=*0.000; 5% FDR adjustment q-value=0.000; Memory/Learning: r=-0.62, *p*=0.001; 5% FDR adjustment q-value=0.008; EF: r=-0.75, *p*=0.000; 5% FDR adjustment q-value=0.000]. APOE4 and Clusterin data maintained no significance (*p*>0.05; data not shown).

#### Predictive capacity of Reelin for AUD cognitive status

Given that only Reelin presented a significant negative correlation with cognition in patients and was the only one with significant differences between patients with vs. without impairment (Student’s t test) (Supplement 5), we determined its predictive capacity by Multiple Linear Regression analyses and found that it could predict cognitive performance in GCF, Memory/Learning, and to a large extent in EF [adjusted R^2^=0.44, *p*=0.000; Memory/Learning: adjusted R^2^=0.36, *p*=0.001; EF: adjusted R^2^=0.53, *p*=0.000]. Reelin was found to predict almost 50% of the performance in general performance (GCF) and more than half (>50%) of the performance found in EF (there was no statistical significance for Visuospatial Cognition (*p*>0.05; data not shown)).

In addition, a Binary Logistic Regression analysis was performed for Reelin to understand the prediction that its concentration had for presenting cognitive impairment by these domains in patients. The model passed the Hosmer-Lemeshow test for all 3 cognitive domains and showed a good calibration [GCF: X^2^_7_=7.92, *p*=0.34; Memory/Learning: X^2^_8_=11.91, *p*=0.15; EF: X^2^_8_=5.69, *p*=0.68]. The results of the prediction analysis have been presented in Table 4. Specifically, Reelin contribute to significantly differentiate between patients with cognitive deterioration vs. those without it in general intelligence (GCF) [Odds Ratio=4.51 (1.37-13.79), *p*=0.13] and EF [Odds Ratio=6.55 (1.68-25.53), *p*=0.07]. The Nagelkerke R^2^ value was: GCF=0.42 and EF=0.54, indicating that Reelin explained 42.3% and more than half (54.0%) of the variance in the dependent variables (impairment in GCF and EF), respectively.

**Table 4.**
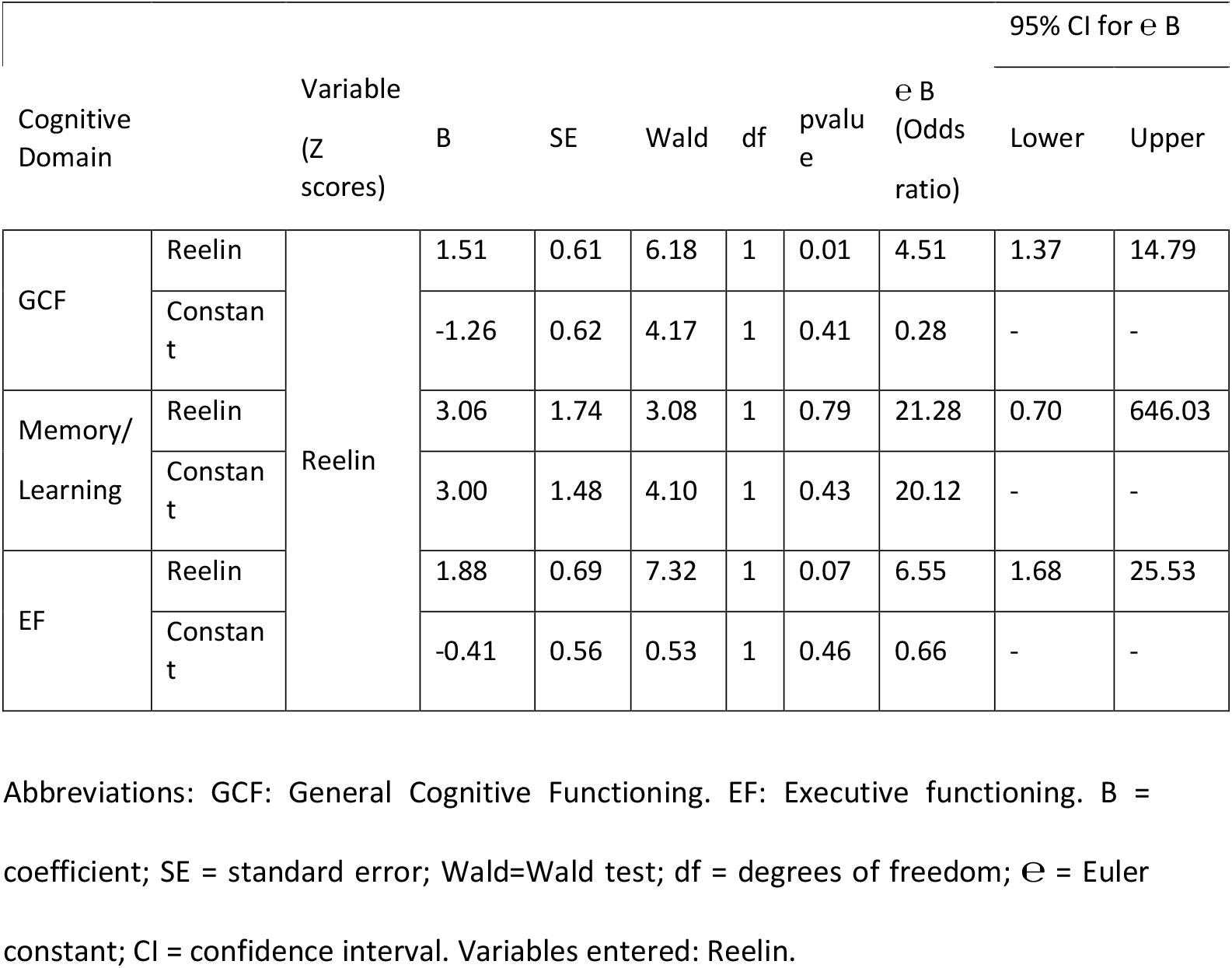
Stepwise logistic regression model based on plasma Reelin concentrations for AUD patients

A ROC curve was performed to predict GCF and EF deficit because plasma Reelin levels were shown to be significant in the Binary Logistic Regression analysis (see ROC curves in Fig. 4 A, B). In concrete, AUC was as follows: [GCF: 0.90, *p*=0.002 (Fig. 4A); EF: 0.88, *p*=0.002 (Fig. 4B)]. The most optimal cut-off point was set at 0.36 (sensitivity: 1; specificity: 0.76) for GCF and 0.36 (sensitivity: 0.90; specificity: 0.86) for EF, according to the maximum values of the Youden index (*J*).

**FIGURE 4.**
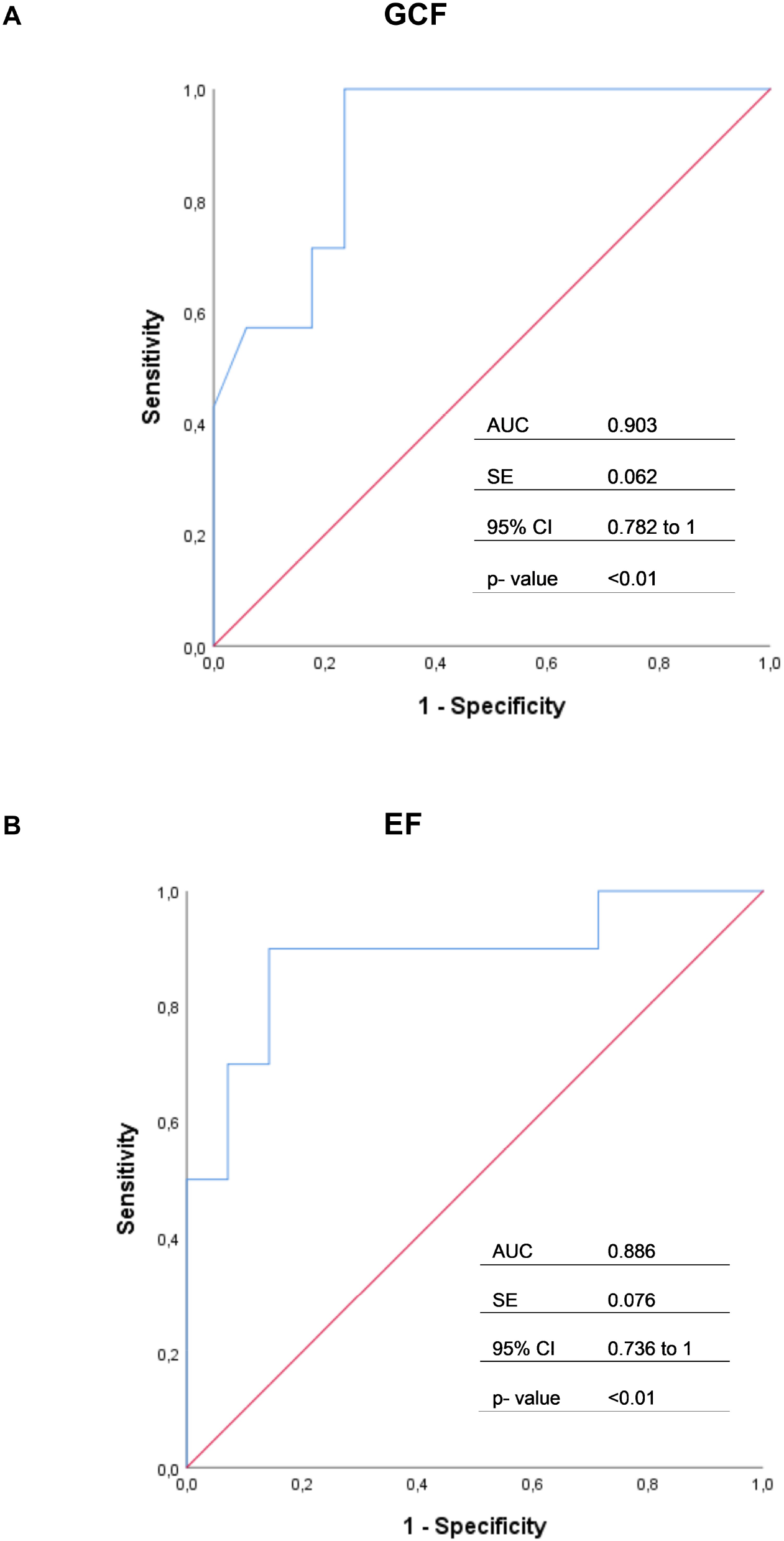
ROC analysis for predictive logistic models of AUD group, using one predictor (Reelin plasma levels), demonstrating the tradeoff between sensitivity (y-axis) and 1-specificity (x-axis). (**A**) GCF (**B**) EF. Both axes of the graph include values between 0 and 1 (0% to 100%). The line drawn from the point 0.0 to the point 1.0 is called the reference diagonal, or the of non-discrimination. Abbreviations: AUC = Area Under the Curve; SE = Standard Error; CI = Confidence Interval.

#### VLDLR and ApoER2 levels in PBMCs and cognition

Significant positive correlations in patients were found between ApoER2 and GFC (r=0.03, *p*<0.05) and Memory/Learning scores (r=0.44, *p*<0.05). Similarly, a significant positive correlation was observed between VLDLR levels and Memory/Learning (r=0.53, p<0.01) and EF (r= 0.44, p<0.05) in patients. No other relations were found neither in correlation analysis, nor in Logistic/Linear Regression (*p*>0.05; data not shown).

## DISCUSSION

One of the genetic risk factors recently found to modulate cognition in alcohol consumption is the presence of the ApoE-ε4 allele (Slayday et al., 2020; Downer et al., 2014; Kim et al., 2012). It is believed that the presence of the ApoE-ε4 allele codes for an aberrant APOE protein, the APOE4, but most of the studies focused on genetic variants with no studies of the real presence of APOE4 in plasma. The development of novel patented methodologies allows the determination of the presence and even the precise quantification of APOE4 protein levels in plasma (Calero et al., 2018). Genetic studies about being ApoE-ε4 carrier and vulnerability to AD and cognition impairment are also widespread (Serrano-Pozo et al., 2021) while scarce investigation has been done regarding APOE4 plasma levels, with a very recent article investigating the pathogenic effect of peripheral APOE4 expression in AD-induced cognitive impairment (Liu et al., 2022). Additionally, most studies in relation to alcohol consumption and general cognition are prospective (Slayday et al., 2020; Downer et al., 2014; Anttila et al., 2004; Dufouil et al., 2000; Reas et al., 2019), with no studies conducted in abstinent AUD patients and no focus on specific cognitive domains. Thus, in the present study we checked plasma APOE4 expression and levels as well as other proteins (Reelin, Clusterin) which APOE4 shares receptors (ApoER2, VLDLR) with, and their relationship with different cognitive domains in a cohort of abstinent AUD patients. Results showed a prominent role for Reelin, which reaches peak plasma levels in patients that express APOE4, predicting cognitive deterioration in AUD.

In line with the previous prospective findings in alcohol consumption, here we found a negative effect of plasma APOE4 presence on cognition (GCF, Memory/Learning and EF) in AUD patients and an interaction between the disorder and APOE4for the Memory/Learning domain. Specifically, AUD patients expressing APOE4 plasma levels may be more vulnerable to the cognitive effects induced by alcohol, being at increased risk for detrimental Memory/Learning performance. Memory is, in fact, the most affected domain in patients, with a very high percentage of deterioration. Although APOE4 has been previously associated with memory loss especially in the context of AD (Najmet al., 2019), our study is the first to report, to our knowledge, its association with memory impairment in AUD-diagnosed patients.

While ApoE-ε4 has been associated with histopathological changes in AD (Emrani et al, 2020), Reelin has been linked to be a protective factor in the maintenance of normal brain function (Levenson et al., 2008). In fact, alteration of Reelin gene in humans is thought to be involved in the pathogenesis of some neuropsychiatric disorders (Ovadia & Shifman, 2011;Wang et al., 2014; Bufill et al., 2015; Li et al., 2015; Fehér et al., 2015). Whereas most studies have linked reduced levels of Reelin both in blood and postmortem brain with neuropsychiatric disorders such as autism (Fatemi et al., 2002, 2005), bipolar disorder (Guidotti et al., 2002; Torrey et al., 2005) and schizophrenia (Guidotti et al., 2002; Impagnatiello et al., 1998; Eastwood and Harrison, 2006), we found a significant increase of Reelin in the plasma of AUD patients compared to controls, in line with other studies. For example, increased expression levels of Reelin in the CSF (Sáez-Valero et al., 2003) and frontal cortex (Botella-López et al., 2006) have been found in AD. Moreover, we observed that the increase in plasma Reelin was much more pronounced when the AUD patients expressed APOE4 plasma levels, meaning that patients expressing the APOE4 isoform were those with the highest levels of Reelin. Interestingly, Reelin was related to and predicted cognitive impairment and cognition in AUD patients, and not in controls, so that patients with plasma APOE4 expression and increased plasma Reelin levels showed higher cognitive impairment, even potentiating the negative effects of the presence of APOE4 itself. In this respect, Reelin negatively correlated with GCF, Memory/Learning and EF and predicted, on one hand, cognitive impairment by domains in GCF and EF and, on the other hand, the degree of cognitive performance in GCF, Memory/Learning and, to a large extent, in EF. So, the cognitive domain most predicted by Reelin in our sample is EF, considered as a complex higher level of mental activities, in line with the literature that links Reelin to higher cognitive function (Ishii et al., 2016).

Reelin deficiency appears to be related to the onset of cognitive deficits (Falconer, 1951; Brigman et at., 2006; Krueger et al., 2006) and, in contrast, here we found that patients with the presence of cognitive deficits are those with the highest Reelin levels. It is very interesting to note that some studies pointed out that the isoform of APOE expressed by the subjects differentially affects the signaling of Reelin. Thus, the APOE4 isoform impairs the receptor recycling to the surface upon activation of Reelin, leading to Reelin resistance to signaling (Lane-Donovan & Herz, 2017). Another described main effect of apoE4 is to impair the NMDA receptor phosphorylation by Reelin, which results in impairments in synaptic plasticity (Chen et al., 2010) and APOE4 can also lower the levels of ApoER2 (Safieh et al., 2019). In this regard, we found lower levels of ApoER2 and VLDL in the PBMCs of patients expressing APOE4, who had also lower scores in cognitive performance, especially for Memory/Learning.

VLDLR and ApoER2 receptors have been identified within the CNS (Lane-Donovan & Herz, 2017) and peripherally (Yakovlev et al., 2012). For example, VLDLR was described as an endothelial cell receptor influencing inflammation (Yakovlev et al., 2012). Little information is available about the presence and function of VLDL and ApoE4 in PBMCs, although both appear to be present in these immune cells (The Human Protein Atlas, https://www.proteinatlas.org/). Here, we detected the presence of both receptors in PBMCs, with lower levels in AUD patients versus controls. Patients expressing the APOE4 isoform had the worst cognitive performance, especially for Memory/Learning. These results agree with studies showing that knock-out mice for these receptors were associated with defects in memory formation and long-term potentiation (Weeber et al., 2002), disruption of neuronal migration (Trommsdorff et al., 1999) or alterations in prepulse inhibition function (Barr et al., 2007), and specially with the Reelin resistance induced by APOE4 explained above.

Clusterin is another apolipoprotein studied here which shares receptors with APOE4 and Reelin (Dlugosz & Nimpf, 2018). Clusterin has been related, among others, to neurodegenerative and cardiovascular diseases, aging, oxidative stress and inflammation (Rodríguez-Rivera et al., 2020; Calero et al., 2000; Trougakos, 2013), as well as changes in cognition and progression to impairment (Thambisetty et al., 2012; Ha et al., 2020; Yang et al., 2019). Here, we have shown that the alterations in Clusterin levels in AUD was unrelated to the cognitive loss. As a whole, we could point to a key role for APOE4 and ApoER2/VLDLR in the cognitive status of AUD patients.

We are aware of the limitations of the same size in this study to extrapolate results to the population. Despite the sample has been age-controlled and sex-matched, validation of results in a larger sample is still needed. Another aspect to be highlight is that the novel eQuant technique does not discriminate homozygous APOE ε4/ε4 from heterozygous APOE ε3/ε4 so we cannot compare these results with genetic studies. However, in the actual context of this study the presence or absence of APOE4 in plasma is a more valuable measure for us that the genetic variants, and the eQuant is an efficient and powerful technique that discriminate specifically the APOE4 from other isoforms such as APOE3.

In conclusion, the presence of APOE4 in plasma has a negative effect on AUD cognition and involves an increase in plasma Reelin levels in these patients. Our data are consistent with a hypothetical model of Reelin resistance and APOE4-induced downregulation of peripheral ApoER2 and VLDL. The APOE/Reelin receptor signaling cascade presents a potential target in clinical trials for novel therapies. An important task for future research will be to use Reelin as a biomarker for cognitive dysfunction in AUD, due to its high predictive capability, to do an appropriate follow-up of AUD patients in outpatient treatment.

## ACKNOWLEDGEMENTS

We thank the nurses Yolanda Guerrero Roldán, Nazaret Sáiz Briones and Pilar de la Cruz González for their invaluable help in this study.

## Data availability statement

Subjects’ personal information is confidential according to the law. Other data are available upon request to the corresponding author.

