## Supplemental Information for "Reelin plasma levels predict cognitive decline in Alcohol Use Disorder: peak levels in patients expressing plasma APOE4 protein"

### SUPPORTING INFORMATION

#### Supplement 1

All patients were under intervention following the hospital's own structured 'Alcohol Programme' (*Hospital 12 de Octubre* (Madrid, Spain)), which has a well-established protocol, running in the clinical practice for more than 20 years, to attend to alcohol-dependent patients. The program is long lasting, since the patients are followed once (or more) per week during 2 years, approximately.

The 'Alcohol Programme' comprises the following two phases:

(1) Detoxification: for patients admitted with acute symptoms, and

(2) Dishabituation: approximately 2 years of program with group and individual therapy. The group sessions comprises the following sub-phases:

- Phase1 (psychoeducation): 12 sessions
- Phase 2 (prevention of relapse): 16 sessions
- Phase 3 (social abilities): 12 sessions
- Phase 4 (preparation for hospital discharge): variable duration

Individual sessions are conducted with the health professionals (psychologists, psychiatrists and nurses) depending on the specific course and needs of each patient. Some participants are sometimes referred to other hospital units, such as gastroenterology, digestive, or others.

#### Supplement 2

AUDIT and MINI are briefly described below:

The 'Alcohol Use Disorders Identification Test' (AUDIT) is a short questionnaire and a self-report screening instrument that explores risk of alcohol drinking, symptoms of dependence and harmful alcohol consumption. If detected risky consumption, the control participant was excluded from the study.

The "Mini International Neuropsychiatric Interview'' (MINI) is a short structured diagnostic interview that explores the main psychiatric disorders of Axis I of DSM-IV (American Psychiatric Association, 1994). If detected the presence of any psychological disorder, the participant was excluded from the study.

On the other hand, the harmful pattern of alcohol consumption in controls was evaluated in the semi-structured interview quantifying the amount of alcohol consumed in months and abstinence time in days (Table S2).

**Table S2.** Semi-structured interview for patients and controls on alcohol consumption

| | |
|---|---|
| Patients | (A) What was the age of first contact? |
| | (B) What is the current abstinence time in weeks/months? |
| | (C) How many bottles, cans or glasses of beer, wine, cocktails (for instance: drinks containing liquor) did you consume per day? |
| | (D) What was the preferred means of consumption? |
| Controls | (A) What is the current abstinence time in days? |
| | (B) How many bottles, cans or glasses of beer, wine, cocktails (i.e. drinks containing liquor) did you consume per month? |

References (Supplement 2)

1. American Psychiatric Association (APA). Diagnostic and statistical manual of mental disorders. 4th ed. American Psychiatric Association; Washington (DC): 1994.

#### Supplement 3

TEDCA ("Test of detection of cognitive impairment in alcoholism”) (Jurado-Barba, 2017) provides a snapshot of cognitive functioning at the time of application including a compendium of 7 tests that assess three specific cognitive functions: Visuospatial Cognition, Memory/Learning and EF (Table S2). TEDCA was administered by a skilled psychologist who had experience and training in neuropsychological test administration. The assessment was established in patients with at least 4 weeks of abstinence (avoiding contamination by residual withdrawal effects or even alcohol intoxication).

**Table S3.** Battery of neuropsychological tests in TEDCA

| Cognitive domain | Domain description | Neuropsychological test |
|---|---|---|
| Visuospatial cognition | Visuo-perceptive, visuo-spatial and visuoconstruction abilities | Rey Complex Figure Test (Copy Condition) (Rey, 1997) |
| | | Bender Visuo-Motor Gestaltic Test (Bender, 2003) |
| Memory/Learning | Ability to store information in the short and long term | Direct and Inverse Digits, Numbers and Letters, and Learning List from the Wechsler Memory Scale (WMS-III) (Wechsler, 2004) |
| EF | Superior cognitive ability. Alternate/divided attention, working memory, planning, organization, problems resolution, abstraction and response inhibition | Trail Making Test B (Reitan, 1985) |
| | | Similarities Test from the Wechsler Adult Intelligence Scale (WAIS-IV) (Wechsler, 2008) |
| | | A Go-No Go task |

References (Supplement 3)

1. Jurado-Barba R, Martínez A, Sion A, Álvarez-Alonso MJ, Robles A, Quinto-Guillen R, Rubio G. Development of a screening test for cognitive impairment in alcoholic population: TEDCA. Actas Esp Psiquiatr. 2017 Sep;45(5):201-17. Epub 2017 Sep 1. PMID: 29044445.
2. Rey A. Test de Copia y Reproducción de Memoria de Figuras Geométricas Complejas. Madrid: TEA Ediciones; 1997.
3. Bender L. Test Guestáltico Visomotor. Buenos Aires: Paidós; 2003.
4. Wechsler D, Pereña J. WMS-III: Escala de memoria de Wechsler III. Madrid: TEA Ediciones; 2004
5. Reitan R, Wolfson D. The Halstead–Reitan Neuropsychological Test Battery: Therapy and clinical interpretation. Tucson, AZ: Neuropsychological Press; 1985.
6. Wechsler D. Wechsler adult intelligence scale–FourthEdition (WAIS–IV). San Antonio, TX: Pearson; 2008.

**Supplement 4**

Reelin, Clusterin, VLDLR and ApoER2 were determined by Human Enzyme-Linked Immunosorbent Assay (ELISA) kits following the manufacturer's instructions, as described below.

Reelin and Clusterin levels in plasma

Plasma Reelin Levels were determined using a commercially available kit (Product #: Human Reln (Reelin) Elisa Kit; reference: HUFI01785; catalog number: EH2121) (Fine test). Plasma samples were 1:2 diluted and the final reaction was measured by optical density (OD) at 450 nm with a spectrophotometer (Molecular Devices®). Reelin concentrations were obtained in pg/mL and expressed in ng/mL. Measurable concentration ranges from 15.625-1000 pg/mL, sensitivity was 9.375 pg/mL intra-assay coefficient of variation was less than 8%.

Plasma Clusterin levels were measured by the ELISA commercially available kit (Product #: RayBio®Human Clusterin Elisa kit; reference: ELH-Clusterin) (RayBiotech). Plasma samples were 1:50000 diluted and with biotinylated anti-human Clusterin antibody and HRP-conjugated streptavidin. The intensity of the color was measured at 450 nm in a spectrophotometer (Molecular Devices®). Results were obtained in pg/mL and expressed in µg/mL. The minimum detectable dose of Human Clusterin was 15pg/mL (Sensitivity) and intra-assay coefficient of variation was less than 10%.

<u>VLDLR and ApoER2 levels in PBMCs</u>

Cellular pellets were resuspended in NP40 lysis buffer (50mM Tris (pH 7.4), 150mM NaCl, 1% NP-40 and 5nM EDTA) (Catalog Number: J60766.AK) with protease and phosphatase inhibitors. Protein concentration was determined by the DC Protein Assay Reagents Package (Bio-Rad Laboratories) (Ref: 5000116) after sonication for 15 s.

VLDLR was assayed by ELISA kit (Product#: Human VLDLR (Very Low-density lipoprotein receptor) Elisa Kit; catalog number: EH1122) (Fine Test). Samples were diluted 1:10.The detection range was from 0.156 to 10 ng/mL, with a sensitivity of 0.094 mL.

ApoER2 was assayed by ELISA kit (Product #: Human LRP8 (Low-density lipoprotein receptor-related protein 8) Elisa kit; catalog number: EH1110) (Fine Test). Samples were diluted 1:2. The detection range of the assay was 0.313 to 20 ng/mL and a sensitivity of 0.188 ng/mL.

For both VLDLR and ApoER2, samples were incubated in microtiter wells with an antibody that only recognized human VLDLR and LRP8, respectively. The OD absorbance was read at 450 nm in Microplate Reader EnSpire (Perkin Elmer). Data were normalized by the amount of total protein.

#### Supplement 5

Patients were classified according to the presence or absence of impairment by each cognitive domain (patients with vs. without impairment). Significant differences were found for Reelin in GCF, Memory/Learning and EF (higher levels in patients with presence of impairment) (Student's t test) [GCF: $t(22)=-3.33$, $p<0.01$; Memory/Learning: $t(22)=-4.16$; EF: $t(22)=-4.26$, $p<0.01$; Visuospatial Cognition: $p>0.05$; data not shown]. APOE4 and Clusterin showed no differences between these groups ($p>0.05$; data not shown).
